# Automated Machine Learning for High-Throughput Image-Based Plant Phenotyping

**DOI:** 10.1101/2020.12.03.410746

**Authors:** Joshua C.O. Koh, German Spangenberg, Surya Kant

## Abstract

Automated machine learning (AutoML) has been heralded as the next wave in artificial intelligence with its promise to deliver high performance end-to-end machine learning pipelines with minimal effort from the user. AutoML with neural architecture search which searches for the best neural network architectures in deep learning has delivered state-of-the-art performance in computer vision tasks such as image classification and object detection. Using wheat lodging assessment with UAV imagery as an example, we compared the performance of an open-source AutoML framework, AutoKeras in image classification and regression tasks to transfer learning using modern convolutional neural network (CNN) architectures pretrained on the ImageNet dataset. For image classification, transfer learning with Xception and DenseNet-201 achieved best classification accuracy of 93.2%, whereas Autokeras had 92.4% accuracy. For image regression, transfer learning with DenseNet-201 had the best performance (R^2^=0.8303, RMSE=9.55, MAE=7.03, MAPE=12.54%), followed closely by AutoKeras (R^2^=0.8273, RMSE=10.65, MAE=8.24, MAPE=13.87%). Interestingly, in both tasks, AutoKeras generated compact CNN models with up to 40-fold faster inference times compared to the pretrained CNNs. The merits and drawbacks of AutoML compared to transfer learning for image-based plant phenotyping are discussed.

## 1 Introduction

High-throughput plant phenotyping (HTP) plays a crucial role in meeting the increasing demand for large-scale plant evaluation in breeding trials and crop management systems (Mir, Reynolds, Pinto, Khan, & Bhat, 2019; Ninomiya, Baret, & Cheng, 2019; Tardieu, Cabrera-Bosquet, Pridmore, & Bennett, 2017). Concurrent with the development of various ground-based and aerial (e.g. unmanned aerial vehicle, UAV) HTP systems is the rise in use of imaging sensors for phenotyping purposes. Sensors for color (RGB), thermal, spectral (multi- and hyperspectral) and 3D (e.g. LiDAR) imaging have been applied extensively for phenotyping applications encompassing plant morphology, physiology, development and postharvest quality (Jiang & Li, 2020; Mir et al., 2019). Consequently, the meteoric rise in big image data arising from HTP systems necessitates the development of efficient image processing and analytical pipelines. Conventional image analysis pipelines typically involve computer vision tasks (e.g. wheat head counting using object detection) which are addressed through the development of signal processing and/or machine learning (ML) algorithms. However, these algorithms are sensitive to image quality (e.g. illumination, sharpness, distortion) variations and do not tend to generalize well across datasets with different imaging conditions (Jiang & Li, 2020). Although traditional ML approaches have to some degree improved upon algorithm generalization, most of them still fall short of the current phenotyping demands and require significant expert guidance in designing features that are invariant to imaging environments. To this end, deep learning (DL), a subset of ML has emerged in recent years as the leading answer to meeting these challenges. One key benefit of DL is that features are automatically learned from the input data, thereby negating the need for laborious manual feature extraction and allow well-generalized models to be trained using datasets from diverse imaging environments. A common DL architecture is deep convolutional neural networks (CNNs) which have delivered state-of-the-art (SOTA) performance for computer vision tasks such as image classification/regression, object detection and image segmentation (Khan, Sohail, Zahoora, & Qureshi, 2020; Minaee et al., 2020; Wu, Sahoo, & Hoi, 2020). The progress of transfer learning, a technique which allows the use of pre-trained SOTA CNNs as base models in DL and the availability of public DL libraries have contributed to the exponential adoption of DL in plant phenotyping. Deep CNN approaches for image-based plant phenotyping have been applied for plant stress evaluation, plant development characterization and crop postharvest quality assessment (Chandra, Desai, Guo, & Balasubramanian, 2020; Jiang & Li, 2020; Watt et al., 2020). However, not all modern CNN solutions can be readily implemented for plant phenotyping applications and adoption will require extra efforts which may be technically challenging (Jiang & Li, 2020). In addition, the process of building a high-performance DL network for a specific task is time-consuming, resource expensive and relies heavily on human expertise through a trial-and-error process (Jiang & Li, 2020; Khan et al., 2020).

Following the exponential growth of computing power and availability of cloud computing resources in recent years, automated machine learning (AutoML) has received tremendous attention in both industry and academia. AutoML provides an attractive alternative to the manual ML practice as it promises to deliver high-performance end-to-end ML pipelines covering data preparation (cleaning and preprocessing), feature engineering (extraction, selection and construction), model generation (selection and hyperparameter tuning) and model evaluation requiring minimal effort or intervention from the user (X. He, Zhao, & Chu, 2019; Truong et al., 2019; Zoller & Huber, 2019). AutoML services have become a standard offering in many technology companies, for example Cloud ML by Google and SageMaker by Amazon. Early work by Zoph et al. (Zoph & Le, 2016) highlighted the potential of AutoML in which a recurrent network was trained with reinforcement learning to automatically search for the best performing architecture. Since then, research interest in AutoML has exploded, with a primary focus on neural architecture search (NAS) which seeks to construct well-performing neural architectures through the selection and combination of various basic modules from a predefined search space (Elsken, Metzen, & Hutter, 2019; X. He et al., 2019; Wistuba, Rawat, & Pedapati, 2019). NAS algorithms can be categorized based on three dimensions: the search space which defines the type of model that can be designed, the search strategy used to explore the search space and the performance estimation strategy which evaluates the performance of possible models from its design (Elsken et al., 2019; Wistuba et al., 2019). NAS-generated models have achieved SOTA performance and outperformed manually designed architectures on computer vision tasks such as image classification (Tan & Le, 2019), object detection (Wang et al., 2019) and semantic segmentation (Weng, Zhou, Li, & Qiu, 2019).

However, despite AutoML showing great promise for computer vision tasks, to the best of our knowledge, no study has used AutoML for image-based plant phenotyping. To address this gap in knowledge, we examined the application of AutoML for image-based plant phenotyping using wheat lodging assessment with UAV imagery as an example. The performance of an open-source AutoML system, AutoKeras was compared to transfer learning using pretrained CNN architectures on image classification and image regression tasks. For image classification, plot images were classified as either non-lodged or lodged; for image regression, lodged plot images were used as inputs to predict lodging scores. The merits and drawbacks of AutoML compared to transfer learning for image-based plant phenotyping are discussed.

## 2 Materials and Methods

### 2.1 Field Experiment

A wheat breeding experiment was conducted at Agriculture Victoria, Horsham, Australia during the winter-spring cropping season of 2018 (Lat:36°44’35.21”S Lon:142°6’18.01”E) (Figure 1). Seeds were sown to a planting density of 150 plants/m^2^ in individual plots measuring 5 m long and 1 m wide (5 m^2^), with a total of 1,248 plots. Severe wind events toward the end of the cropping season (30^th^ November – 9^th^ December) resulted in significant lodging of wheat plots across the experiment. Ground truth labels for “lodged” and “non-lodged” were provided by an experienced field technician and a plant scientist.

**Figure 1.**
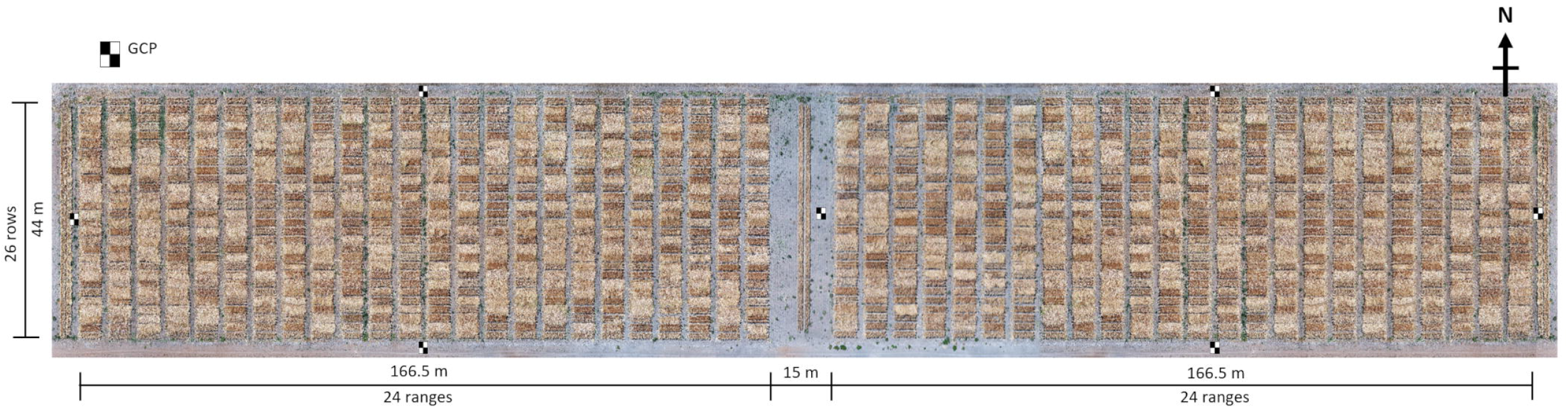
Wheat breeding field experiment. Ground control points (GCPs) indicated in figure.

### 2.2 Image Acquisition and Processing

High resolution aerial imaging of wheat plots with different lodging grades was performed on 11^th^ December 2018. Aerial imagery was acquired on a DJI Matrice M600 Pro (Shenzhen DJI Sciences and Technologies Ltd., China) UAV with a Sony A7RIII RGB camera (35.9 mm x 24.0 mm sensor size, 42.4 megapixels resolution) mounted on a DJI Ronin MX gimbal. Flight planning and automatic mission control was performed using DJI’s iOS application Ground Station Pro (GS Pro). The camera was equipped with a 55 mm fixed focal length lens and set to 1 s interval shooting with JPEG format in shutter priority mode. Images were geotagged using a GeoSnap Express system (Field of View, USA). The flight mission was performed at an altitude of 45 m with front and side overlap of 75% under clear sky conditions. Seven black and white checkered square panels (38 cm x 38 cm) were distributed in the field experiment to serve as ground control points (GCPs) for accurate geo-positioning of images (Figure 1). A real-time kinematic global positioning system (RTK-GPS) receiver, EMLID Reach RS (https://emlid.com) was used to record the centre of each panel with < 1 centimeter accuracy.

Images were imported into Pix4Dmapper version 4.4.12 (Pix4D, Switzerland) to generate an orthomosaic image, with the coordinates of the GCPs used for geo-rectification. The resulting orthomosaic had a ground sampling distance (GSD) of 0.32 cm/pixel. Individual plot images were clipped and saved in TIFF format from the orthomosaic using a field plot map with polygons corresponding to the experimental plot dimension of 5 m x 1 m in ArcGIS Pro version 2.5.1 (Esri, USA).

### 2.3 Lodging Assessment

A two-stage assessment of lodging was performed in this study and the results were used as the basis for image classification and image regression tasks (see Section 2.4) (Figure 2). The image classification task corresponded to the first stage of assessment in which the lodging status i.e. whether the plot is lodged (yes) or non-lodged (no) was provided by the ground truth data and this could be verified easily from visual inspection of the high-resolution plot images (Figure 3). The image regression task corresponded to the second stage of assessment where plots identified as lodged were evaluated using a modified lodging score based on the method of Fisher and Stapper (Fischer & Stapper, 1987):

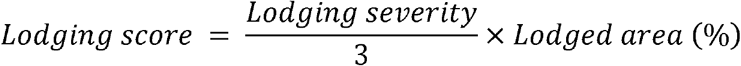

**Figure 2.**
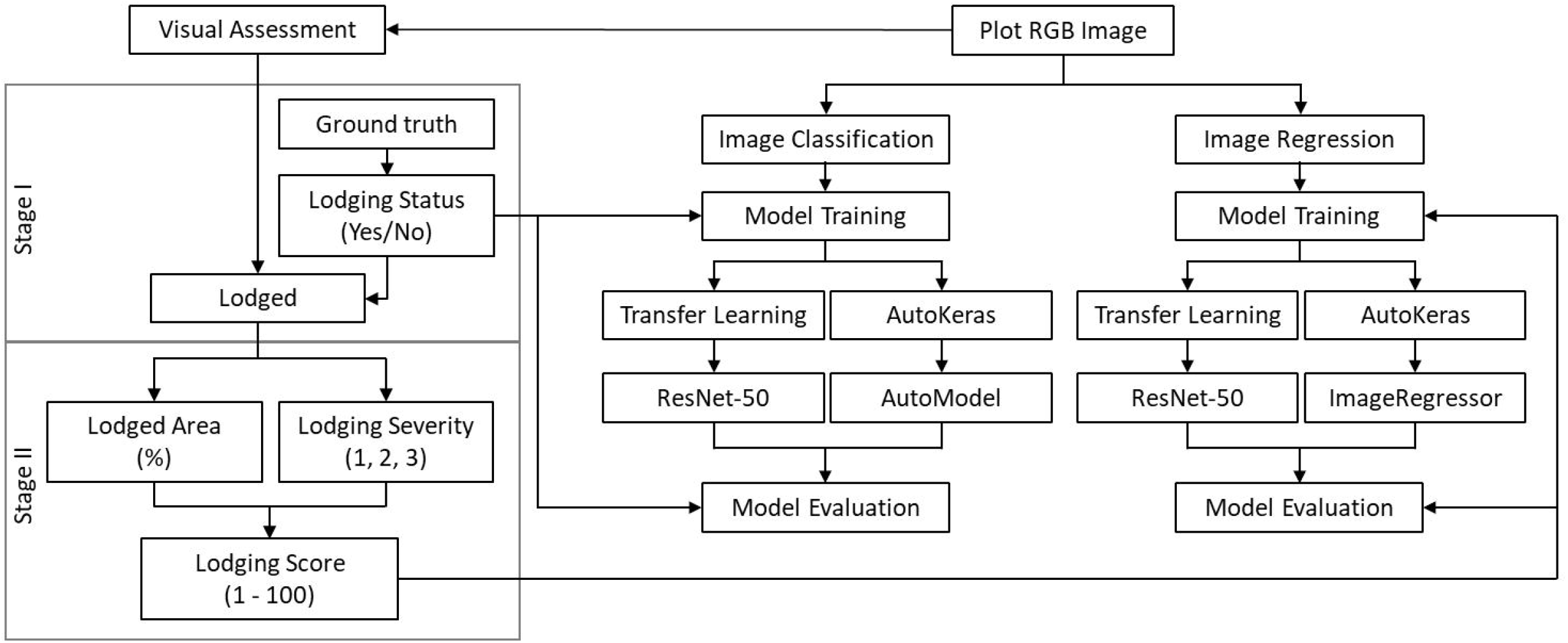
Experimental workflow for wheat lodging assessement.

**Figure 3.**
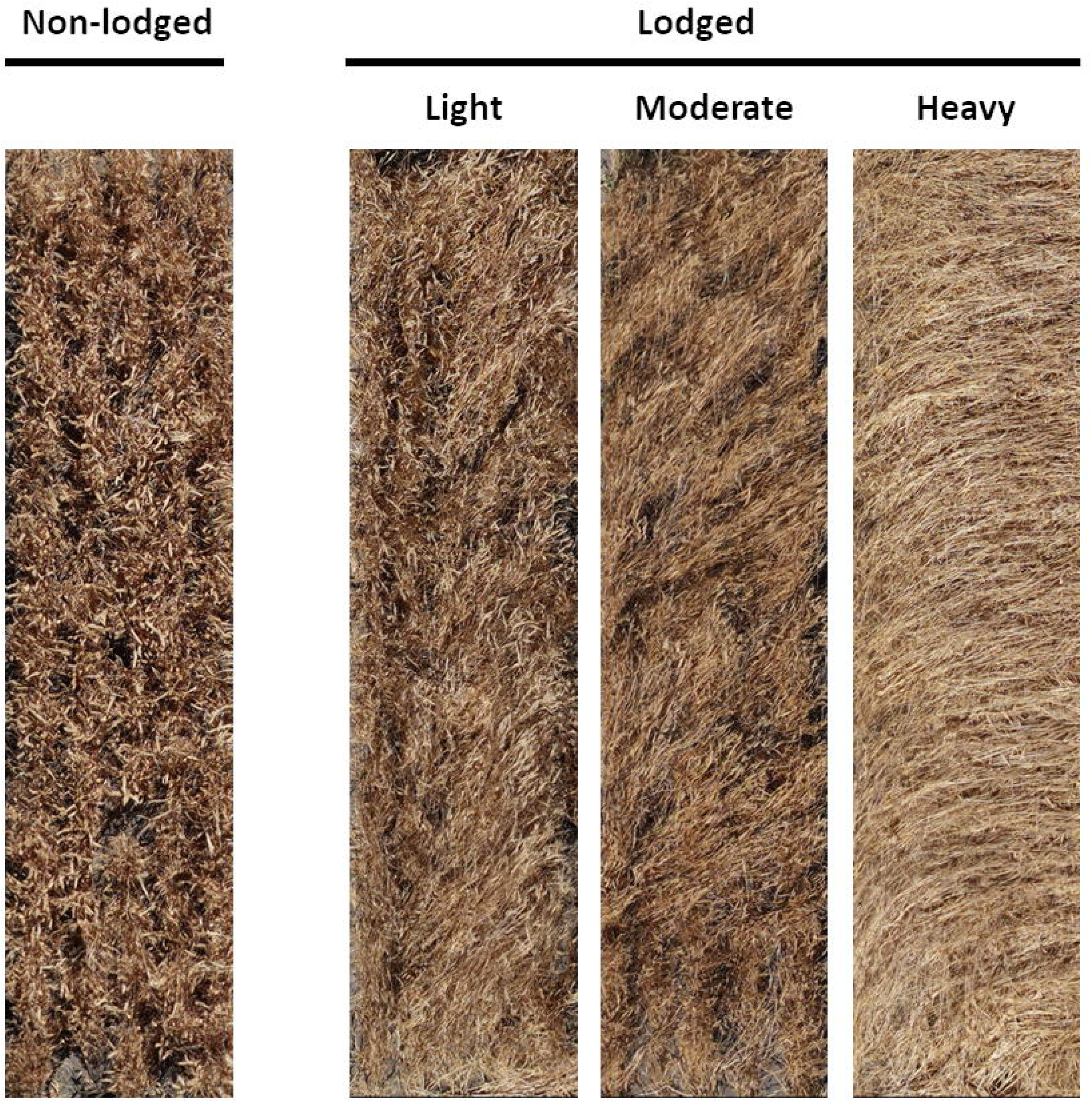
Different wheat lodging severities. Wheat plot images were first classified as non-lodged or lodged using ground truth data. Plots identified as lodged were assessed visually and divided into three lodging severities (light, moderate and heavy) based on lodging angles.

Lodging severity values of 1 to 3 were assigned to three main grades of lodging based on the inclination angle between the wheat plant and the vertical line as follows: light lodging (severity 1; 10° – 30°), moderate lodging (severity 2; 30° – 60°) and heavy lodging (severity 3; 60° – 90°)(Figure 3)(Sun et al., 2019). The lodged area (%) was determined visually from the plot images as the percentage of area lodged in the plot in proportion to the total plot area. The derived lodging score ranged between values of 1 – 100, with a score of 100 indicating that the entire plot was lodged with heavy lodging.

### 2.4 Deep Learning Experiments

Deep learning experiments were conducted in the Python 3.7 environment. Performance of the open-source AutoML framework, AutoKeras (Jin, Song, & Hu, 2019) version 1.0.1 in image classification and image regression was compared to a manual approach using transfer learning with pretrained modern CNN architectures implemented in Keras, Tensorflow-GPU version 2.1. For the image classification task, a binary classification scheme assigning individual plot images to either non-lodged (class 0) or lodged (class 1) was performed as the images had a relatively balanced distribution between the two classes. For the image regression task, plot images were used as inputs to predict the lodging score. Training and evaluation of the models were performed on a NVIDIA Titan RTX GPU (24 GB of GPU memory) at SmartSense iHub, Agriculture Victoria.

#### 2.4.1 Training, Validation and Test Datasets

For image classification, the image dataset consisted of 1,248 plot images with 528 plots identified as non-lodged (class 0) and 720 plots identified as lodged (class 1). Images were first resized to the dimensions of 128 width x 128 height x 3 channels (Section 2.4.2) and these were split 80:20 (seed number=123) into training (998 images) and test (250 images) datasets. For image regression, the 720 resized plot images identified as lodged were split 80:20 (seed number=123) into training (576 images) and test (144 images) datasets. Images were fed directly into AutoKeras without preprocessing as this was done automatically by AutoKeras. In contrast, images were preprocessed to the ResNet-50 format using the provided preprocess_input function in Keras. For model training on both image classification and regression, the training dataset was split further 80:20 (seed number=456) into training and validation datasets. The validation dataset is used to evaluate training efficacy, with lower validation loss (as defined by the loss function, Section 2.4.2) indicating a better trained model. Performance of trained models was evaluated on the test dataset (section 2.2.4).

#### 2.4.2 AutoML with AutoKeras

AutoKeras is an open-source AutoML framework built using Keras which implements state-of-the-art NAS algorithms for computer vision and machine learning tasks (Jin et al., 2019). It is also the only open-source NAS framework to offer both image classification and regression abilities at the time of this study. In our study, we experienced great difficulty in getting AutoKeras to stably complete experiments in default settings due to errors relating to GPU memory usage and model tuning. This is not entirely unexpected as beginning version 1.0, AutoKeras has undergone significant application programming interface (API) and system architecture redesign to incorporate KerasTuner ver. 1.0 and Tensorflow ver. 2.0. This change was necessary for AutoKeras to capitalize on recent developments in NAS and the DL framework, Tensorflow, in addition to providing support for the latest graphics processing unit (GPU) hardware. To partly circumvent the existing issues, we had to implement two approaches for the DL experiments to stably complete up to 100 trials, which is the number of models evaluated by AutoKeras (i.e. 100 trials = 100 models). Firstly, all images were resized to the dimensions of 128 x 128 x 3 and secondly, for image classification, we had to implement a custom image classifier using the provided AutoModel class in AutoKeras. We were not successful in completing experiments beyond 100 trials, as such only results up to 100 trials were presented in this study.

For image classification, a custom image classifier was defined using the AutoModel class which allows the user to define a custom model by connecting modules/blocks in AutoKeras (Figure 4). In most cases, the user only needs to define the input node(s) and output head(s) of the AutoModel, as the rest is inferred by AutoModel itself. In our case, the input nodes were first an ImageInput class accepting image inputs (128 x 128 x 3), which in turn was connected to an ImageBlock class which selects iteratively from either a ResNet (K. He, Zhang, Ren, & Sun, 2016), ResNext (Xie, Girshick, Dollár, Tu, & He, 2017), Xception (Chollet, 2017) or simple CNN building blocks to construct neural networks of varying complexity and depth. The input nodes were joined to a single output head, the ClassificationHead class which performed the binary classification (Figure 4). The AutoModel was fitted to the training dataset with the tuner set as “bayesian”, loss function as “binary_crossentropy”, evaluation metrics as “accuracy” and 200 training epochs (rounds) for 10, 25, 50 and 100 trials with a seed number of 10. For image regression, the default Autokeras image regression class, ImageRegressor was fitted to the training dataset with the loss function as mean squared error (MSE), evaluation metrics as mean absolute error (MAE) and mean absolute percentage error (MAPE), and 200 training epochs for 10, 25, 50 and 100 trials with a seed number of 45 (Figure 4).

**Figure 4.**
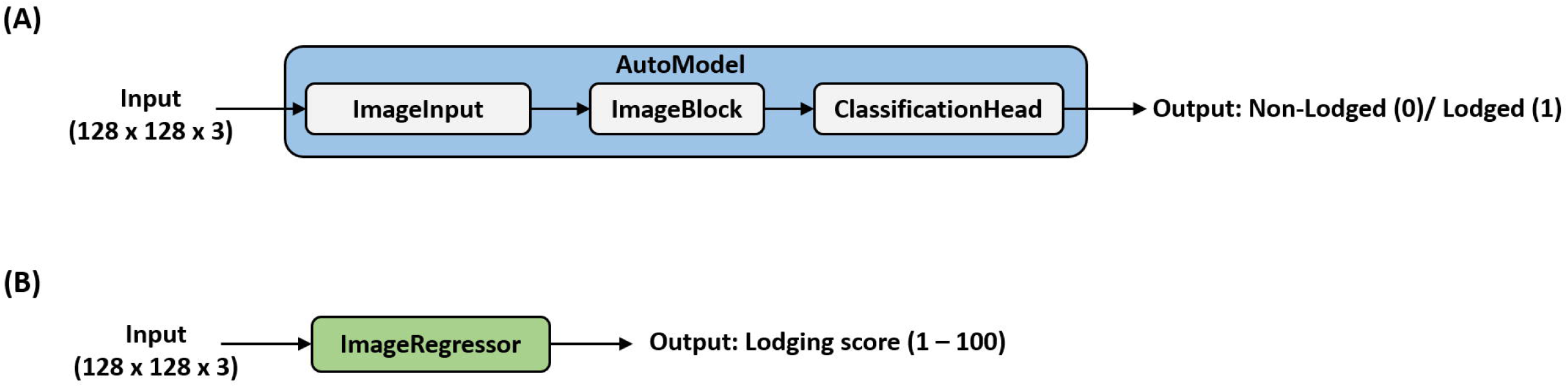
Automated machine learning with AutoKeras. (A) AutoModel for image classification. (B) ImageRegressor for image regression.

The performance of the best models from 10, 25, 50 and 100 trials were evaluated on their respective test datasets (Section 2.4.4) and exported as Keras models to allow neural network visualization using the open-source tool, Netron (https://github.com/lutzroeder/netron).

#### 2.4.3 Transfer Learning with pretrained CNNs

Transfer learning is a popular approach in DL where a pretrained model is reused as the starting point for a model on a second task (Jiang & Li, 2020). This allows the user to rapidly deploy complex neural networks, including state-of-the-art DL architectures without incurring time and computing costs. In this study, transfer learning was performed using VGG networks (Simonyan & Zisserman, 2015), residual networks (ResNets)(K. He et al., 2016), InceptionV3(Szegedy, Vanhoucke, Ioffe, Shlens, & Wojna, 2016), Xception (Chollet, 2017) and densely connected CNNs (DenseNets)(Huang, Liu, & Weinberger, 2017) pretrained on the ImageNet dataset. These networks were implemented in Keras as a base model using the provided Keras API with the following parameters: weights=“imagenet”, include_top=False and input_shape=(128, 128, 3) (Figure 5). Output from the base model was joined to a global average pooling 2D layer and connected to a final dense layer, with the activation function set as either “sigmoid” for image classification or “linear” for image regression. The model was compiled with the batch size as 32, optimizer as ‘Adam’ and corresponding loss functions and evaluation metrics as described in section 2.4.2. Model training occurred in two stages for both image classification and regression tasks: in the first stage (100 epochs), weights of the pre-trained layers were frozen and the Adam optimizer had a higher learning rate (1 × 10^−1^ or 1 × 10^−2^) to allow faster training of the top layers; in the second stage (200 epochs), weights of the pre-trained layers were unfrozen and the Adam optimizer had a smaller learning rate (1 × 10^−2^ to 1 × 10^−5^) to allow fine-tuning of the model. Learning rates were optimized for each pretrained CNN and the values which provided the best model performance are provided in supplementary Table S1. Performance of the trained models was evaluated on their respective test datasets (Section 2.4.4).

**Figure 5.**
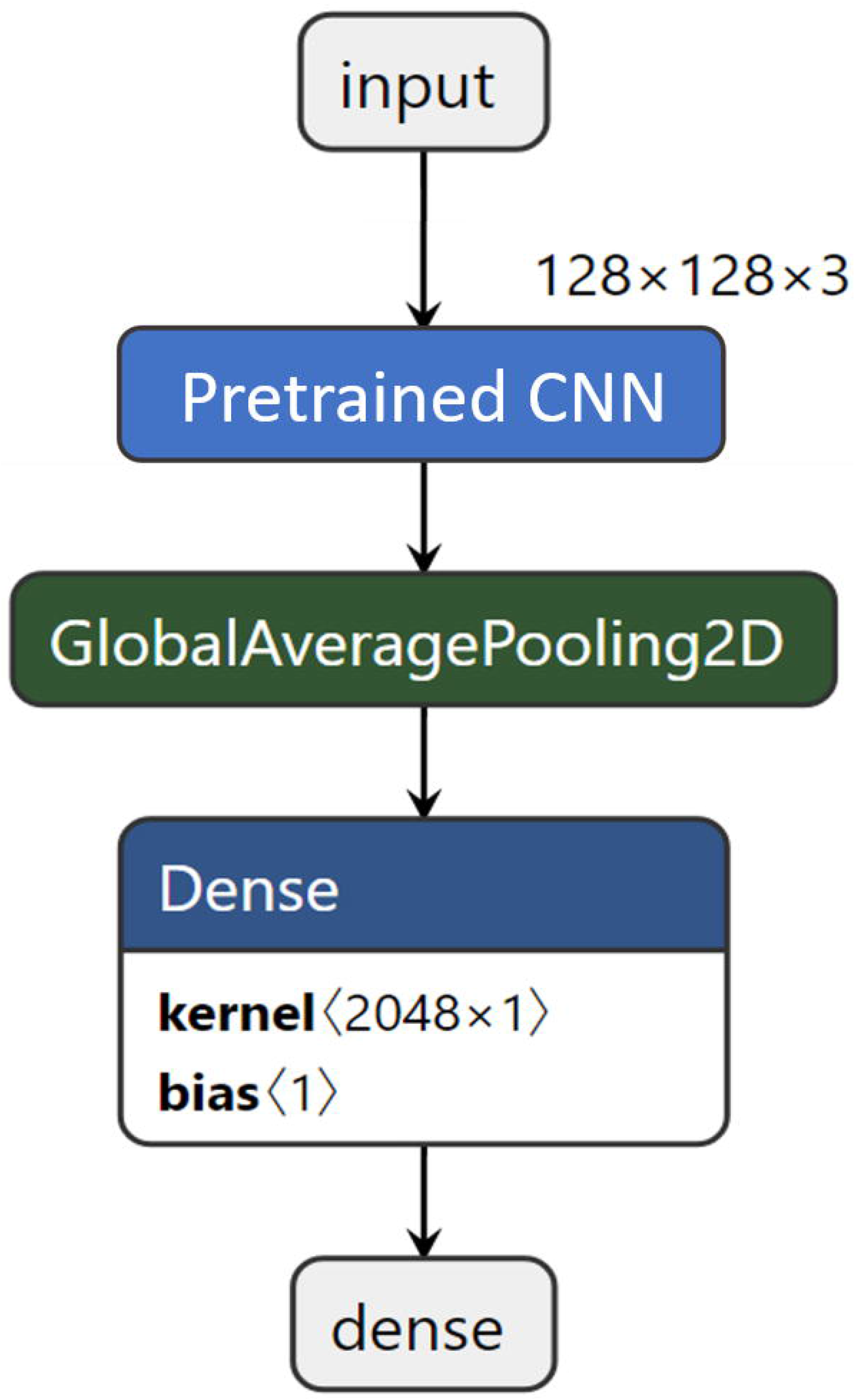
Transfer learning with pretrained CNN architectures. Output from a pretrained CNN was joined to a global average pooling 2D layer and connected to a final dense layer, with the activation function set as either “sigmoid” for image classification or “linear” for image regression.

#### 2.4.4 Model Evaluation Metrics

For image classification, model performance on the test dataset was evaluated using classification accuracy (%). For image regression, in addition to the mean absolute error (MAE) and the mean absolute percentage error (MAPE) provided by AutoKeras and Keras, the coefficient of determination (R^2^) and the root mean-squared error (RMSE) were also calculated to determine model performance on the test dataset. Models were also evaluated based on total model training time (in minutes, min) and inference time on the test dataset presented as mean ± standard deviation in milliseconds (ms).

## 3 Results

### 3.1 Image Classification

Both transfer learning with pretrained CNNs and AutoKeras performed strongly in the image classification task (Table 1). Transfer learning performance with pretrained CNNs ranged from 91.6 to 93.2% classification accuracy, with Xception and DenseNet-201 giving the best overall accuracy of 93.2% (Table 1). Among the pretrained CNNs, InceptionV3 had the fastest training (5.42 min) and inference (100.54 ± 15.08 ms) times, whereas DenseNet-201 had the slowest training (11.79 min) and inference (188.09 ± 24.19 ms) times. In comparison, AutoKeras (AK) performance ranged from 86.8 to 92.4% accuracy, with performance improving as more models (trials) were evaluated (Table 1). The best AutoKeras model was discovered from 100 trials and had the same 92.4% accuracy as the ResNet-50 (Table 1). Impressively, AutoKeras was able to achieve this result using a simple 2-layers CNN (43,859 parameters) consisting only of a single 2D convolutional layer (Figure 6) as opposed to the 50-layers deep ResNet-50 architecture (~23.6 million parameters). Incidentally, the 2-layers CNN had the fastest overall inference time (5.71 ± 0.12 ms) on the test dataset compared to other models, which was ~18-fold faster compared to the InceptionV3 and up to ~33-fold faster compared to the DenseNet-201 (Table 1). However, model training times for AutoKeras were significantly higher compared to the transfer learning approaches, with the longest training time of 251 min recorded for 100 trials, which was ~21-fold higher compared to the DenseNet-201 (Table 1). Examination of the model architectures returned by AutoKeras revealed that the best model architecture resulting from the 10 and 25 trials was a deep CNN model comparable in depth and complexity to the ResNet-50 (Figure S1), highlighting the ability of AutoKeras to explore deep CNN architectures even in a small model search space. Subsequently, when the model search space was extended to 50 and 100 trials, the best model architecture discovered by AutoKeras was the 2-layers CNN model (Figure 6).

**Table 1.** Model performance metrics for image classification.

| Network | Parameters | Training (min) | Inference (ms) | Accuracy (%) |
| --- | --- | --- | --- | --- |
| VGG16 | 14,715,201 | 6.03 | 146.70 ± 20.54 | 92.0 |
| VGG19 | 20,024,897 | 7.01 | 161.70 ± 25.87 | 91.6 |
| ResNet-50 | 23,589,761 | 5.89 | 119.41 ± 15.52 | 92.4 |
| ResNet-101 | 42,660,225 | 9.88 | 186.73 ± 26.14 | 92.8 |
| InceptionV3 | 21,804,833 | 5.42 | 100.54 ± 15.08 | 92.8 |
| Xception | 20,863,529 | 9.06 | 148.20 ± 20.75 | 93.2 |
| DenseNet-169 | 12,644,545 | 9.23 | 152.83 ± 22.93 | 92.8 |
| DenseNet-201 | 18,323,905 | 11.79 | 188.09 ± 24.19 | 93.2 |
| AK-10_trials | 23,566,856 | 16.06 | 102.34 ± 14.33 | 86.8 |
| AK-25_trials | 23,566,856 | 29.18 | 110.45 ± 13.25 | 88.4 |
| AK-50_trials | 43,859 | 102.43 | 5.83 ± 0.64 | 89.6 |
| AK-100_trials | 43,859 | 251.80 | 5.71 ± 0.12 | 92.4 |

**Figure 6.**
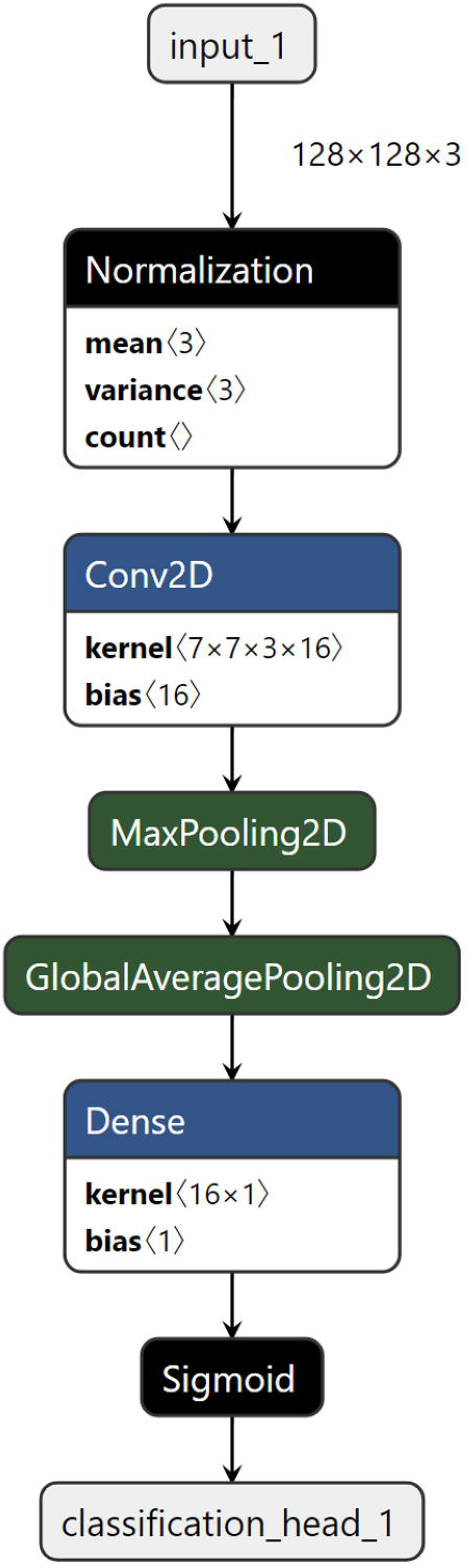
Best AutoKeras model architecture for image classification. A simple 2-layers CNN model with 43,895 parameters was the best architecture discovered in 50 and 100 trials. The best AutoKeras classification performance was provided by the model from 100 trials.

### 3.2 Image Regression

For the image regression task, transfer learning with DenseNet-201 gave the best overall performance (R^2^=0.8303, RMSE=9.55, MAE=7.03, MAPE=12.54%), followed closely by AutoKeras (AK) with the model from 100 trials (R^2^=0.8273, RMSE=10.65, MAE=8.24, MAPE=13.87%)(Table 2). The CNN models varied in regression performance, with R^2^ ranging between 0.76 – 0.83. Within the pretrained CNNs, DenseNet-201 had the slowest model training (7.01 min) and inference (117.23 ± 15.25 ms) times, with ResNet-50 having the fastest training (3.55 min) and inference (79.23 ± 10.31 ms) times. For AutoKeras, performance generally improved from 10 to 100 trials (Table 2). AutoKeras was able to achieve the second-best performance using an 8-layers CNN resembling a truncated mini Xception network with 207,560 total parameters (Figure 7). Two prominent features of the original 71-layers deep Xception network, namely the use of depthwise separable convolution layers and skip connections were evident in the AutoKeras model (Figure 7). Notably, the mini Xception network outperformed the original pretrained Xception network (R^2^=0.7709, RMSE=11.08, MAE=8.22, MAPE=13.51%)(Table 2). Not surprisingly, the mini Xception network had the fastest inference time (2.87 ± 0.12 ms) compared to the other models, which was ~27-fold faster compared to the ResNet-50 and up to 41-fold faster compared to the DenseNet-201 (Table 2). However, model training times for AutoKeras was again significantly higher compared to the transfer learning approaches, with the longest training time of 325 min recorded for 100 trials, which was ~46 fold higher compared to the DenseNet-201 (Table 2). Examination of the model architectures returned by AutoKeras revealed that the best model architecture resulting from the 10 and 25 trials was a deep CNN model (Figure S2) whereas the best model architecture discovered from 50 and 100 trials was the 8-layers mini Xception model (Figure 7).

**Table 2.** Model performance metrics for image regression.

| Network | Parameters | Training (min) | Inference (ms) | R<sup>2</sup> | RMSE | MAE | MAPE (%) |
| --- | --- | --- | --- | --- | --- | --- | --- |
| VGG16 | 14,715,201 | 3.71 | 90.86 ± 12.72 | 0.7590 | 11.37 | 8.97 | 14.02 |
| VGG19 | 20,024,897 | 4.32 | 103.86 ± 16.43 | 0.7707 | 11.03 | 9.19 | 16.01 |
| ResNet-50 | 23,589,761 | 3.55 | 79.23 ± 10.31 | 0.7844 | 10.79 | 8.28 | 15.51 |
| ResNet-101 | 42,660,225 | 5.85 | 114.85 ± 16.08 | 0.7730 | 11.10 | 8.38 | 15.67 |
| InceptionV3 | 21,804,833 | 3.32 | 62.18 ± 9.33 | 0.7642 | 11.09 | 8.07 | 13.90 |
| Xception | 20,863,529 | 5.33 | 92.91 ± 13.01 | 0.7709 | 11.08 | 8.22 | 13.51 |
| DenseNet-169 | 12,644,545 | 6.65 | 94.25 ± 14.14 | 0.7985 | 10.31 | 7.68 | 13.63 |
| DenseNet-201 | 18,323,905 | 7.01 | 117.23 ± 15.25 | 0.8303 | 9.55 | 7.03 | 12.54 |
| AK-10_trials | 23,566,856 | 32.25 | 80.26 ± 0.14 | 0.7568 | 12.43 | 9.54 | 14.55 |
| AK-25_trials | 23,566,856 | 123.08 | $82.36 \pm 0.12$ | 0.7772 | 12.28 | 8.62 | 14.38 |
| AK-50_trials | 207,560 | 184.91 | $2.86 \pm 0.11$ | 0.8133 | 10.71 | 8.31 | 13.92 |
| AK-100_trials | 207,560 | 325.62 | $2.87 \pm 0.12$ | 0.8273 | 10.65 | 8.24 | 13.87 |

**Figure 7.**
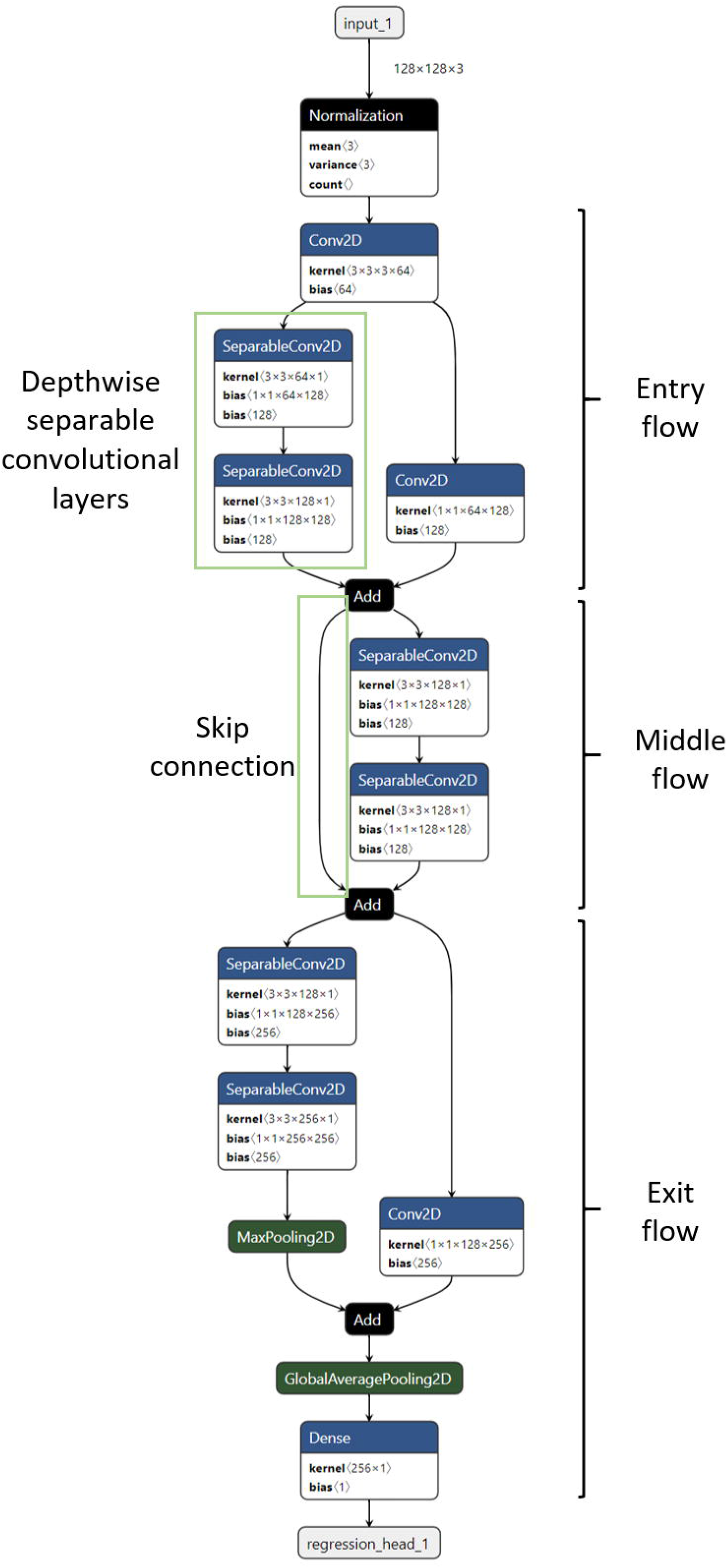
Best AutoKeras model architecture for image regression. An 8-layers mini Xception model with 207,560 parameters was the best architecture discovered in 50 and 100 trials. Three main parts (entry, middle and exit flows) and two key features (examples indicated by green boxes), namely the depthwise separable convolutional layers and skip connections from the original Xception network (Chollet, 2017) can be discerned from the architecture. The best AutoKeras regression performance was provided by the model from 100 trials.

## 4 Discussion

Using wheat lodging assessment as an example, we compared the performance of an open-source AutoML framework, AutoKeras to transfer learning using modern CNN architectures for image classification and image regression. As a testament to the power and efficacy of modern DL architectures for computer vision tasks, both AutoKeras and transfer learning approaches performed well in this study, with transfer learning exhibiting a slight performance advantage over AutoKeras.

For the image classification task, plot images were classified as either non-lodged or lodged. The best classification performance of 93.2% was jointly achieved by transfer learning with Xception and DenseNet-201 networks. This is not entirely surprising as both Xception (Chollet, 2017) and DenseNet (Huang et al., 2017) were developed later (chronologically speaking) as improved architectures compared to the other CNNs in this study. In contrast, the best AutoKeras model (from 100 trials) achieved an accuracy of 92.4%, which is the same as those obtained by transfer learning with ResNet-50. Classification results in this study are comparable to those reported in a smaller study which classified 465 UAV-acquired wheat plot multispectral images (red, green, blue channels used in models) as either non-lodged/lodged with a hand-crafted deep neural network, LodgedNet (97.7% accuracy) and other modern DL networks (97.7 – 100.0% accuracies) via transfer learning (Mardanisamani et al., 2019). The higher classification accuracies reported in that study may be due to the use of image data augmentation (i.e. increasing training dataset via image transformations) which generally improves model performance and that images were acquired pre-maturity (plants were still green) allowing variations in color to potentially contribute more informative features for modelling (Mardanisamani et al., 2019; Sun et al., 2019). Results in our study suggest that NAS generated models can provide competitive performance compared to modern human designed CNN architectures, with transfer learning using pretrained CNNs exhibiting a slight performance advantage (~1% improvement) over AutoML models. A recent survey comparing the performance of manually designed models against those generated by NAS algorithms on the CIFAR-10 dataset, a public dataset commonly used for benchmarking in image classification, found that the top two best-performing models were both manually designed models (X. He et al., 2019). However, the gap between the manual and AutoML models was very small (<1% accuracy difference). In our study, AutoKeras was able to achieve results comparable to those of the 50-layers deep ResNet-50 model (~25 million parameters) using only a simple 2-layers CNN model (43,859 parameters). The 2-layers CNN had the fastest inference time (5.71 ± 0.12 ms) compared to the other models, which was up to 33-fold faster compared to the DenseNet-201 which had the slowest inference time. As such, the 2-layers CNN could prove useful for real-time inferencing, but it’s simple or shallow architecture raises concern about its generalizability across different datasets. This can be addressed in future studies by using datasets derived from multiple trials or imaging conditions for model training to obtain a solution generalized across different environments.

For the image regression task, lodged plot images were used as inputs to predict the lodging score. The best performance (R^2^=0.8303, RMSE=9.55, MAE=7.03, MAPE=12.54%) was obtained using transfer learning with DenseNet-201, followed closely by AutoKeras (R^2^=0.8273, RMSE=10.65, MAE=8.24, MAPE=13.87%) with the model discovered from 100 trials. In both image classification and regression tasks, transfer learning with DenseNet-201 achieved the best results. DenseNet can be considered as an evolved version of the ResNet (K. He et al., 2016), where the outputs of the previous layers are merged via concatenation with succeeding layers to form blocks of densely connected layers (Huang et al., 2017). However, similarly for image classification, the DenseNet-201 had the slowest inference time (117.23 ± 15.25 ms) on the test dataset in image regression, making it less suitable for time-critical applications such as real-time inferencing. In comparison, the AutoKeras model resembled a mini 8-layers Xception model (207,560 parameters) and had the fastest inference time (2.87 ± 0.12 ms) on the test dataset, which was ~41-fold faster compared to the DenseNet-201. In its original form, the Xception network is 71-layers deep (~23 million parameters) and consists of three parts: the entry flow, middle flow and exit flow (Chollet, 2017). These three parts and two key features of the Xception network, namely the depthwise separable convolutions and skip connections originally proposed in ResNet were discernable from the mini Xception model. Research in the area of network pruning which compresses deep neural network through the removal/pruning of redundant parameters showed that it is possible to have equally performant models with up to 97% of the parameters pruned (Salama, Ostapenko, Klein, & Nabi, 2019). Although dissimilar to network pruning, as evidenced in our study, AutoKeras can discover efficient and compact model architectures through the NAS process. However, NAS generated models are typically limited to variants or combinations of modules derived from existing human designed CNN architectures (Elsken et al., 2019; X. He et al., 2019; Wistuba et al., 2019); although recent innovations in NAS have uncovered novel CNN architectures such as SpineNet for object detection (Du et al., 2019).

The lodging score originally proposed by Fisher and Stapper (Fischer & Stapper, 1987) is calculated from the lodging angle and lodged area (lodging score = angle of lodging from vertical position/90 x % lodged area). In our study, the angle of lodging is replaced by lodging severity, which is a score of 1 to 3 assigned to light, moderate and heavy lodging grades as determined by visual assessment of lodged plot images. Consequently, under- or over-estimation of lodging scores may happen for plots within the same lodging severity grade. For example, plots with 100% lodged area and lodging angle of 65^0^(lodging severity 3) and 90^0^ (lodging severity 3) will have the same lodging score of 100 in this study as opposed to scores of 72 and 100 according to the original method. This may partly account for the model prediction errors in the image regression task. Nonetheless, the modified lodging score allowed a rapid evaluation of wheat lodging based on visual assessments of UAV imagery and was useful as a target for image regression. For detailed assessment of lodging and model performance in lodging score prediction, future studies should incorporate manual ground-truthing of the lodging angle and lodged area to enable a more accurate calculation of lodging scores.

One of the challenges in this study was getting AutoKeras to perform stably and complete the DL experiments. Initial attempts were often met with an out of memory (OOM) error message, arising from AutoKeras trying to load models too large to fit in the GPU memory. Prior to version 1.0, AutoKeras had a GPU memory adaptation function which limits the size of neural networks according to the GPU memory (Jin et al., 2019). However, beginning version 1.0, this function is no longer implemented in AutoKeras (personal communication with Haifeng Jin, author of AutoKeras) and may partially account for the OOM errors. To partly circumvent this issue, we had to resize all input images (1544 × 371) to a smaller size of 128 × 128, which allowed AutoKeras to complete experiments up to 100 trials. The impact of the downsized images on AutoKeras model performance would need to be ascertained in future studies, although the 128 × 128 image size is within common ranges observed in DL models, for example, LodgedNet (64 × 128)(Mardanisamani et al., 2019) and established modern CNN architectures (224 × 224)(K. He et al., 2016; Huang et al., 2017; Simonyan & Zisserman, 2015). As AutoKeras is undergoing active development, we are hopeful that issues encountered in our study will be resolved in future releases. It will be interesting to explore AutoKeras’ performance with larger model search space (>100 trials) using higher resolution input images coupled with image data augmentation where appropriate in future studies.

Transfer learning using existing modern CNN architectures achieved better results compared to AutoML in both image classification and regression tasks in this study. However, a major limitation is that these CNN models have been trained using 3-channels RGB images and this prevents direct application of these models for image sources beyond 3-channels, such as multispectral and hyperspectral images (Mardanisamani et al., 2019). In addition, existing CNN architectures may not always provide the best performance compared to custom-designed models. However, modification of existing CNN architectures or manually designing CNN models are time-consuming and technically challenging. In this regard, AutoML provides an attractive alternative as it can deliver CNN models with good performance out-of-the-box and can accept inputs of varying sizes and dimensions, making it ideal for use on diverse sensor-derived data including multispectral and hyperspectral imagery. Furthermore, an added benefit of AutoML is its potential through NAS to discover compact model architectures which are ideal for real-time inferencing. However, to ensure generalizability of the NAS-generated models across datasets, it is vital to employ training dataset representative of the diverse trial environments and imaging conditions in future studies. The primary downside of AutoML is the long training times (hours as opposed to minutes) required to achieve models with competitive performance compared to transfer learning even when using modern GPU hardware. The time and GPU computational costs associated with AutoML hinders it from being widely adopted by DL practitioners for now. However, this is expected to be offset in time by the rapid growth in GPU computing power and concurrent rise in GPU affordability. Another concern relating to AutoML is the reproducibility of results owing to the stochastic nature of NAS (Li & Talwalkar, 2019). This concern can largely be addressed through making available all datasets, source codes (including exact seeds used) and best models reported in the NAS study – a practice embraced in this study (see Data Availability).

Results in our study demonstrate that transfer learning with modern CNNs performed better compared to AutoML, although the performance differences were minimal. For most computer vision tasks using RGB image datasets, transfer learning with existing CNNs will provide a good starting point and should yield satisfactory results in most cases with minimal effort and time. However, for plant phenotyping applications that are time-critical and generate image datasets beyond the standard RGB 3-channels, AutoML is a good alternative to manual DL approaches and should be in the toolbox of both novice and expert users alike. For field-based crop phenotyping, portable multispectral and hyperspectral sensors are becoming common on ground-based and aerial HTP platforms (Mir et al., 2019), providing ample avenues for AutoML application. Moving forward, with the exponential rise in GPU computing power and strong interests in NAS research, AutoML systems are expected to become more ubiquitous. In tandem with existing DL practices, they can contribute significantly towards streamlined development of image analytical pipelines for HTP systems integral to improving breeding program and crop management efficiencies. Results in our study provide a basis for the adoption and application of AutoML systems for high-throughput image-based plant phenotyping.

## 5 Acknowledgments

We thank Dennis Ward and Emily Thoday-Kennedy for technical support in conducting the field experiment.

## 6 Author Contributions

JCOK designed the deep learning experiments and analyzed the data. JCOK, GS and SK conceived and designed the study. SK managed the field experiment. JCOK, GS and SK wrote and edited the paper. All authors read and approved the final manuscript.

## 7 Funding

This study is funded by Agriculture Victoria.

## 8 Conflict of Interest Statement

The authors declare that the research was conducted in the absence of any commercial or financial relationships that could be construed as a potential conflict of interest.

## 9 Data Availability

Dataset containing 1,248 digital images (1,544 × 371 pixels) of individual wheat plots with varying grades of lodging and corresponding lodging data are available on Zenodo (Joshua Koh, German Spangenberg, & Surya Kant, 2020). Ground truth data for lodging status and calculated lodging scores are provided as a CSV file. Source codes required to replicate the analyses in this article and the best performing models reported for AutoKeras are provided in a GitHub repository (Joshua Koh, German Spangenberg, & Surya Kant, 2020).

## 11 Supplementary Materials

**Figure S1. Best AutoKeras model architecture from 10 and 25 trials for image classification.**

**Figure S2. Best AutoKeras model architecture from 10 and 25 trials for image regression.**

**Table S1. Adam optimizer learning rates for pretrained CNNs fine-tuning.**

## Notes

https://github.com/AVR-SmartSense/automl_paper

https://zenodo.org/record/3952422#.X8lmUbNxUuU

### Competing Interest Statement

The authors have declared no competing interest.

