## Supplementary figures and images for "Automated Machine Learning for High-Throughput Image-Based Plant Phenotyping"

### Figure S1

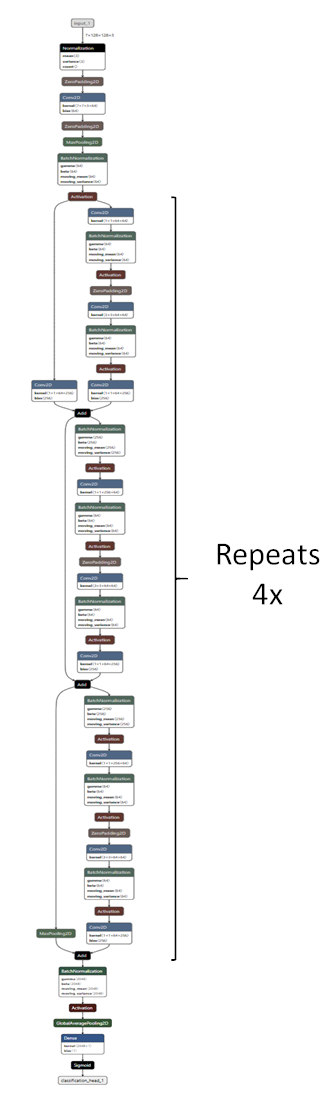

### Figure S2

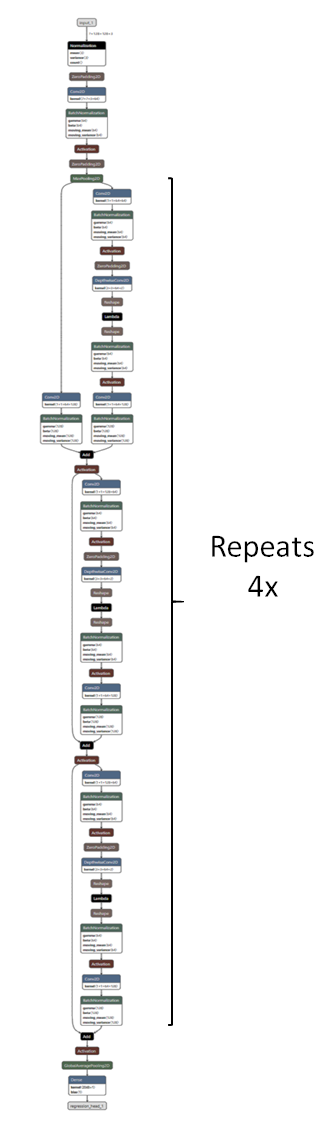
